# *In vitro* Isolation of *Treponema pallidum* subsp. *pallidum* from Fresh and Frozen Needle Aspirates of Primary Experimental Syphilis Lesions

**DOI:** 10.1101/2022.09.13.507848

**Authors:** Lauren C. Tantalo, Barbara J. Molini, Mahashweta Bose, Jeffrey D. Klausner, Lorenzo Giacani

## Abstract

Isolation of *Treponema pallidum* subsp. *pallidum* strains still relies on rabbit intratesticular inoculation of clinical samples. Here, we report an alternative isolation approach based on the inoculation of fresh and frozen needle aspirates of primary experimental lesions into culture plates suitable for *in vitro* propagation of the syphilis agent.

**SUMMARY:** We report on a rabbit-free *in vitro* isolation approach for syphilis strains that can be easily adapted to clinical specimens.

## INTRODUCTION

In 1912, Nichols and Hough reported the successful isolation of a strain of *Treponema pallidum* subsp. *pallidum* (*T. pallidum*) using intratesticular inoculation of a brown-and-white rabbit with a darkfield-negative sample of cerebrospinal fluid (CSF) from a syphilis patient (1). The authors concluded that the rabbit model was suitable for inoculation of specimens with low numbers of spirochetes (1). Over a century later, strain isolation through rabbit intratesticular injection still provides investigators with novel strains to study syphilis pathogenesis and vaccine development. The rabbit-dependent procedure for isolation of *T. pallidum* strains can be performed on samples as diverse as blood, lesion exudates, CSF, kidney and liver biopsies, amniotic fluid, and neonatal serum (2-6). Furthermore, recent work demonstrated the rabbit-based isolation of new strains from cryopreserved lesion exudates (7), reducing the need for clinical collection sites to be located near a research facility capable of housing rabbits in a BSL2 environment.

Strain isolation using rabbits, however, has several drawbacks. First, this approach carries a significant financial burden due to the cost of animals, their husbandry, veterinary care, and the need for trained personnel for rabbit monitoring post-inoculation. Second, immune pressure from the rabbit may result in pathogen loss due to immune clearance if strain harvest is not performed timely when the rabbit develops orchitis or, in the absence of it, immediately after seroconversion. Lastly, reports like that of Tong *et al*. (8), who found a low yield of positive results after intratesticular rabbit inoculation. might discourage investigators from adopting rabbit-based strain isolation.

An alternative to sample inoculation into rabbit is offered by the cell culture system suitable for continuous *in vitro* propagation of *T. pallidum* described by Edmondson *et al*. (9, 10). This system has yet to be applied to *T. pallidum* isolation from clinical samples. Here, we report how *in vitro* isolation of syphilis strains can be achieved using needle aspirates from primary experimental lesions. This approach can be easily translated to patient lesion samples to attain rabbit-free isolation of syphilis strains.

## METHODS and RESULTS

To obtain primary lesions, the UW211B *T. pallidum* strain, isolated in 2003 by Christina Marra (11), was revived from a cryopreserved stock and passaged *in vitro* as previously described (12) until rabbit infection. Two male New Zealand White rabbits were inoculated intradermally (ID) on their shaved backs on several sites by injecting 10^6^ *T. pallidum* cells/site. Rabbit backs were kept shaven, and injection sites were monitored daily for lesion development. For the UW211B strain, lesion progression to induration occurs on average at 24 days post-inoculation. Needle aspirates were obtained from indurated lesions that had not yet ulcerated. All animal procedures were approved by the University of Washington Institutional Animal Care & Use Committee (IACUC protocol #4243-01; PI: Lorenzo Giacani), and care was provided per instructions of the Guide for the Care and Use of Laboratory Animals.

On the day preceding lesion aspiration, sterile 24-well culture plates were seeded with Sf1Ep cells following the *T. pallidum in vitro* propagation protocol (13). Three hours before aspirate collection, Sf1Ep cell culture media was replaced with 2 ml of fresh TpCM2 media that had been made the previous day and equilibrated overnight in a Heracell 150i tri-gas incubator (1.5% O_2_, 3.5% CO_2_, 95% N_2_; ThermoFisher, Waltham, MA) at 34°C.

In experiment 1, four lesions were aspirated from one rabbit using a tuberculin syringe with a 25-gauge needle. Following aspiration, the needle/syringe was rinsed with 75 µl of fresh TpCM2 media to recover spirochetes. The concentration of *T. pallidum* cells in suspension was determined by darkfield microscopy (DFM) as described previously (12). Suspensions were then serially diluted in fresh TpCM2 to achieve concentrations ranging from 1.16×10^7^ to 500 *T. pallidum* cells/ml, which provided a subset of samples where spirochetes could not be detected by DFM. Twenty-five microliters of these dilutions were inoculated into plate wells in duplicate. The smallest inoculum corresponded to 12.5 *T. pallidum* cells/well, while the highest was 2.9×10^5^ cells/well.

Plates were incubated for two weeks at 34°C in the tri-gas incubator, with a TpCM2 media exchange at one week. After two weeks, exhausted media was discarded, and cells were sub-cultured following trypsinization, as reported previously (13). At each passage, 18 µl of the culture supernate obtained from each well were used for DFM quantification to assess treponemal growth. The remainder (∼400 µl) was inoculated into a well of a freshly prepared cell culture plate to be passaged again after two additional weeks. This procedure was carried out for a total of 8 weeks.

The results of experiment 1 (Fig.1A), reported as treponemal growth curves determined by DFM count, showed that *T. pallidum* cells could be retrieved from all samples by week 8 post-inoculation, including those samples where *T. pallidum* could no longer be quantified by DFM at the initial timepoint (dashed line on Fig.1A-E). Most of these DFM-negative samples required a blind passage at week 2 post-inoculation (Fig.1A). In this experiment, the smallest inoculum from which *T. pallidum* was reisolated was of 12.5 *T. pallidum* cells, resulting from inoculating 25 µl of a 500 *T. pallidum* cells/ml suspension.

**Figure 1.**
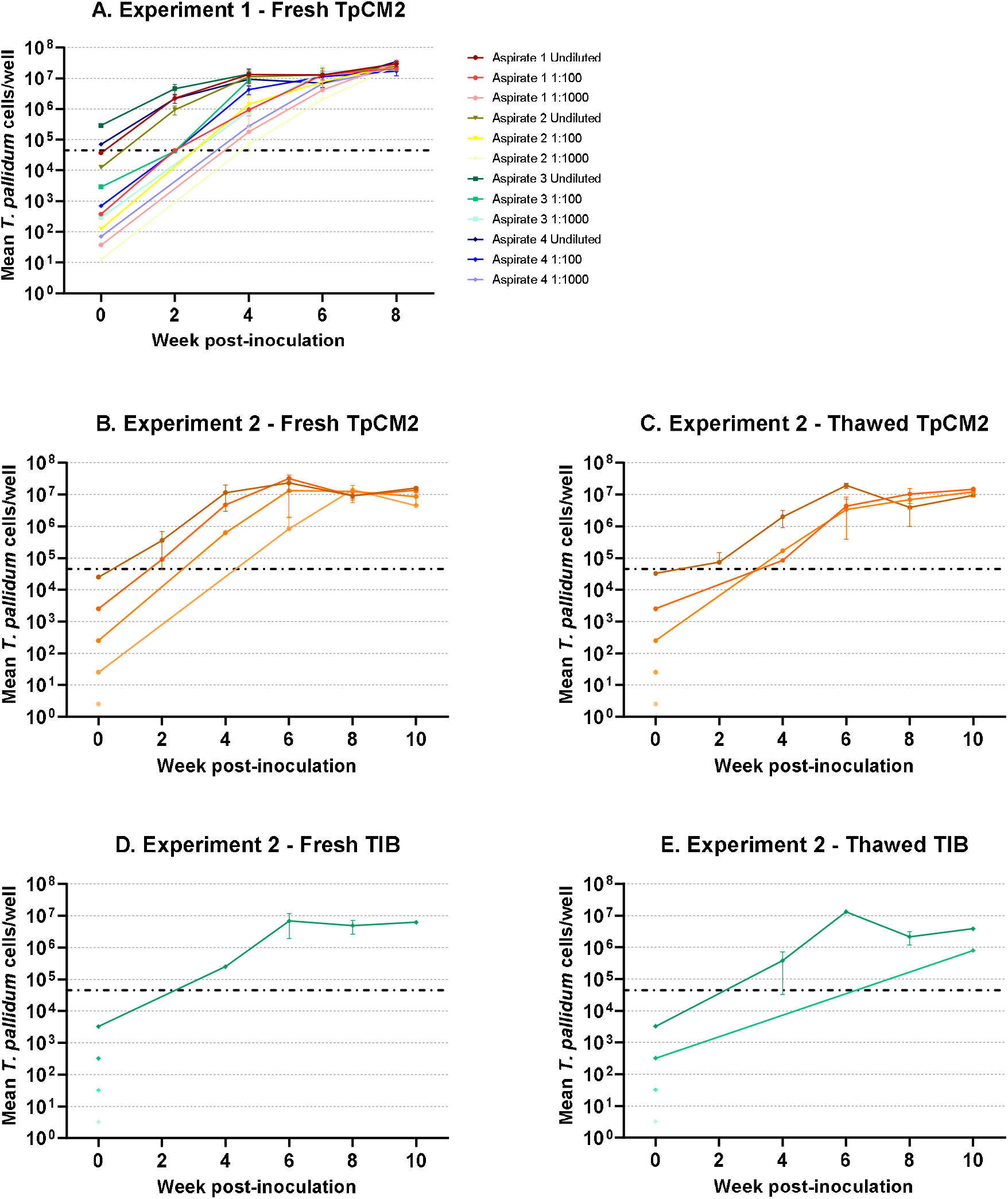
*T. pallidum* growth curves resulting from inoculating samples from lesion aspirates into culture plates suitable for *T. pallidum in vitro* propagation. Lesion aspirates were prepared in TpCM2 (panel A/B), TpCM2 supplemented with glycerol, frozen for 1 week before thawing and subsequent inoculation (panel B), fresh TIB (panel D), and TIB, frozen for 1 week before thawing and subsequent inoculation (panel E). All samples were tested in duplicate. Values indicate mean *T. pallidum* cells/well +/-SD, determined using DFM quantification. Treponemal inoculum at week 0 was calculated based on initial treponemal concentration in the undiluted sample and dilution factor applied. The dashed line represents the DFM limit of detection.

The impact of freezing and thawing on the recovery of *T. pallidum* from aspirates was evaluated in experiment 2. Additionally, this experiment assessed the possibility of replacing TpCM2 with Treponemal Isolation Buffer (TIB; 25% sterile saline, 25% heath-inactivated normal rabbit serum, and 50% sterile glycerol), routinely used in our laboratory to collect and cryopreserve viable treponemes from patient samples and significantly easier to prepare compared to TpCM2. For this second experiment, 2 needle aspirates were obtained from two distinct indurated lesions from the infected rabbit. One aspirate was resuspended and serially diluted in a suitable volume of TpCM2. Half of the volume from these dilutions was directly inoculated into culture plates. To the remaining half, an equal volume of sterile glycerol was added. Samples were stored at -80°C for 1 week, thawed and used to inoculate duplicate wells of a culture plate. The second aspirate was also resuspended in TpCM2 (as TIB high viscosity prevents the release of treponemes from the needle/syringe), but dilutions were performed using TIB (already containing glycerol) instead of TpCM2. Half of the resulting volume was inoculated immediately, and half was cryopreserved for a week prior to inoculation. In these experiments, the minimum inoculum was 2.5 *T. pallidum* cells/well for samples resuspended in TpCM2, and 3.25 *T. pallidum* cells/well for samples in TIB. Sub-culturing was performed as above but for up to 10 weeks post-inoculation.

Results (Fig.1B-E) suggest that treponemal isolation is facilitated both by using TpCM2 buffer and avoiding a freezing step, and by extending the isolation period to 10 weeks. When using fresh TpCM2, treponemes could be reisolated from all inoculated wells but the ones starting with only 2.5 *T. pallidum* cells/well (Fig.1B) by week 6 post-inoculation (Fig.1B). As seen before (Fig.1A) several blind passages were needed to recover spirochetes from wells inoculated with 250 *T. pallidum* cells of fewer. Thawed TpCM2 samples (Fig.1C) yielded a positive result starting with an inoculum of 250 *T. pallidum* cells, but not lower. Treponemes resuspended in fresh TIB (Fig.1D) could be reisolated only when the inoculum consisted of 3,250 cells total, but not lower. Although treponemes could be reisolated from an inoculum of 325 treponemes (Fig.1E) when using thawed TIB, only one of the two wells yielded results and required 4 blind passages before treponemes could be detected.

## DISCUSSION

Albeit preliminary, the present study suggests that properly collected samples from a syphilis patient could be effectively used to isolate a new *T. pallidum* strain using the *in vitro* cultivation system for this spirochete instead of rabbit intratesticular inoculation. For labs already culturing *T. pallidum*, strain isolation requires minimal additional time and labor compared to rabbit-based isolation, while labs already performing tissue culture work can implement this technique with relatively low effort. Soon, we aim to apply this procedure to clinical specimens, such as aspirates of secondary disseminated lesions collected after surface disinfection to avoid sample contamination with the skin flora. *T. pallidum* cultivation is generally performed in the absence of antibiotics, which makes it prone to the growth of bacteria other than *T. pallidum* if sample contamination occurs at collection or proper sterile technique is not applied during routine passages. Although Phosphomycin or Amphotericin B will not harm *T. pallidum* and may be used individually to minimize the chances of losing a sample to contamination (9, 10), these compounds might not always be effective against all contaminants. Our study also suggests that TpCM2 might be the medium of choice for sample collection. This media could be prepared fresh, equilibrated, and stored frozen (9, 10) until use at the collection site. Lastly, additional studies to expand the range of samples that can be used with this procedure, such as secondary lesion biopsies, blood, plasma, CSF, or amniotic fluid, would further encourage the use of this technique to replace rabbit isolation.

